# Epistasis and destabilizing mutations shape gene expression variability in humans via distinct modes of action

**DOI:** 10.1101/026393

**Authors:** Ence Yang, Gang Wang, Jizhou Yang, Beiyan Zhou, Yanan Tian, James J. Cai

## Abstract

Increasing evidence shows that, like phenotypic mean, phenotypic variance is also genetically determined, but the underlying mechanisms of genetic control over the variance remain obscure. Here, we conducted variance-association mapping analyses to identify expression variability QTLs (evQTLs), i.e., genomic loci associated with gene expression variance, in humans. We discovered that common genetic variations may contribute to increasing gene expression variability via two distinct modes of action—epistasis and destabilization. Specifically, the epistasis model explains a quarter of the identified evQTLs, of which the formation is attributed to the presence of “third-party” eQTLs that influence the level of gene expression in a fraction, rather than the entire set, of sampled individuals. The destabilization model explains the other three-quarters of evQTLs, which tend to be associated with mutations that disrupt the stability of the transcription process of genes. To show the destabilizing effect, we measured discordant gene expression between monozygotic twins, and time-course stability of gene expression in single samples using repetitive qRT-PCR assays. The destabilizing evQTL SNPs were found to be associated with more pronounced expression discordance between twin pairs and less stable gene expression in single samples. Together, our results suggest that common SNPs may work interactively or independently to shape the variability of gene expression in humans. These findings contribute to the understanding of the mechanisms of genetic control over phenotypic variance and may have implications for the development of variability-centered analytic methods for quantitative trait mapping.

## Introduction

Quantitative genetics assumes that phenotypic variation—the difference in phenotypic *mean* between individuals—is genetically controlled (1). Phenotypic variation is thus typically referred to as the difference in phenotypic mean among genotypes. This assumption, however, has been challenged. Some recent studies have shown that phenotypic *variance* is also genetically controlled and that the variance itself is a quantitative trait (2–14). It is clear that research on the genetics of phenotypic variance deserves more attention. Understanding how phenotypic variance is controlled is of great importance not only for quantitative genetics but also for evolutionary biology, agricultural and animal sciences, and medicine (5, 11, 15, 16). For example, a greater phenotypic variance may offer more adaptive solutions in evolution (17–19), and thus, genetic factors resulting in more variable phenotypes may become favored as they enable a population to respond more effectively to environmental changes (20–23). In medicine, disease states may emerge when the relevant phenotype of affected individuals goes beyond a threshold. Thus, more variable genotypes will produce a larger proportion of individuals exceeding that threshold than will less variable genotypes, even if these genotypes have the same mean. Therefore, by ignoring the effect of genotypes on phenotypic variance, an important axis of genetic variation contributing to phenotypic differences among individuals has been overlooked (1, 24). In this regard, the lack of empirical studies has hindered the discovery of variance-associated mutations that may contribute to human health-related traits, including those modulating disease susceptibility.

Previous studies have shown the existence of substantial gene expression variability in humans, including significant differences in the magnitude of gene expression variance between groups (25–27). Yet, our understanding of how genetic factors control or modulate gene expression variance remains limited. Promising new developments along this line have come from recent findings in complex trait analysis of gene expression variance (9, 11, 12). Using variance association mapping, we and others have identified genetic loci associated with gene expression variance, called expression variability quantitative trait locus (evQTLs) (11, 12), also known as v-eQTL (9). How evQTLs are originated is not completely known. Epistasis has been widely accepted as a mechanism that introduces phenotypic variability through genetic interactions. In this study, we seek a non-epistatic, more straightforward, explanation—that is, evQTL SNPs (evSNPs) disrupt or destabilize the genetic architecture that buffers stochastic variation in gene expression. We call this explanation the “destabilization model,” which emphasizes the destabilizing effect of a mutation that pushes the gene expression trait out of homeostasis or equilibrium to become less robust. We show that the formation of evQTLs can be explained by using either the epistasis model or the destabilization model. We anticipate that our findings will lay a foundation for developing a new analytical framework that focuses on the contributions of genetic variation to phenotypic variance.

## Results

### Widespread evQTLs in the human genome

We obtained the expression data of 15,124 protein-coding genes measured in 462 lymphoblastoid cell lines (LCLs) by the Geuvadis Project (28). We also obtained the genotype data of 2,885,326 polymorphic sites determined in the same cell lines by the 1,000 Genomes Project (29). After data processing, 326 LCL samples from unrelated individuals of European descent (EUR) were retained for this study (**Materials and Methods**). To identify evQTLs, we first applied the method based on the double generalized linear model (DGLM) (30), which has been previously adopted by us (11, 12) and others (5). In the interest of computational time, we restricted the use of this method in the identification of *cis*-acting evQTLs, so that, for each gene, only those SNPs located within a 1-Mb region of up- and downstream of the transcription start site were considered. On average, each gene has 1,803 SNPs in its *cis*-regions. Using a conservative Bonferroni correction cutoff *P*=1.75×10^-9^ (=0.05/28,494,473), we identified a total of 17,949 *cis*-evQTLs in 1,304 unique genes, accounting for 8.6% of all genes tested (**Figure 1A**) (**Supplementary Table S1**). We repeated this analysis using raw expression data (i.e., the un-normalized RPKM values) and found that more than half (56.7%) of *cis*-evQTLs could be recovered, suggesting that the normalization of data did not profoundly impact the evQTL detection. Next, we adopted a computationally efficient method based on the squared residual value linear model (SVLM) (9, 31) to identify both *cis*- and *trans*-evQTLs. The SVLM method is a two-stage approach: the effect of a tested SNP on gene expression mean (i.e., eQTL effect) is first regressed out, and then the correlation between the squared expression residuals and the SNP genotypes is tested. We applied the SVLM method to all SNPs and tested against all genes. Such an all-against-all strategy, without pre-filtering SNPs by their location relative to tested genes, allowed a systematic detection of *cis*- and *trans*-evQTLs across the entire genome. We used the Benjamini–Hochberg procedure (32) to determine the *P*-value cutoff of 3×10^-9^ to reach the false-discovery rate (FDR) of 10%. At this level, we identified 505 *cis*-evQTLs in 33 unique genes, and 1,008 *trans*-evQTLs in 235 unique genes (**Figure 1B**) (**Supplementary Table S2**). The distribution of *cis*- and *trans*-evQTLs across autosomes (**Figure 1C**) does not show any enrichment of *cis*-evQTLs compared to *trans*-evQTLs. Furthermore, there is a marked discrepancy between the number of *cis*-evQTLs detected using SVLM and DGLM. But, 463 (91.7%) of 505 *cis*-evQTLs detected by SVLM were also detected by DGLM (**Supplementary Table S1**), suggesting the discrepancy was due to the lack of power of the SVLM method. Indeed, our simulation-based study showed that the SVLM method had only half of the power of DGLM at the sample size of 300 (**Supplementary Fig. S1**). The discrepancy may also be attributed to the greater multiple testing burden associated with applying SVLM in the all-against-all tests.

**Fig. 1.**
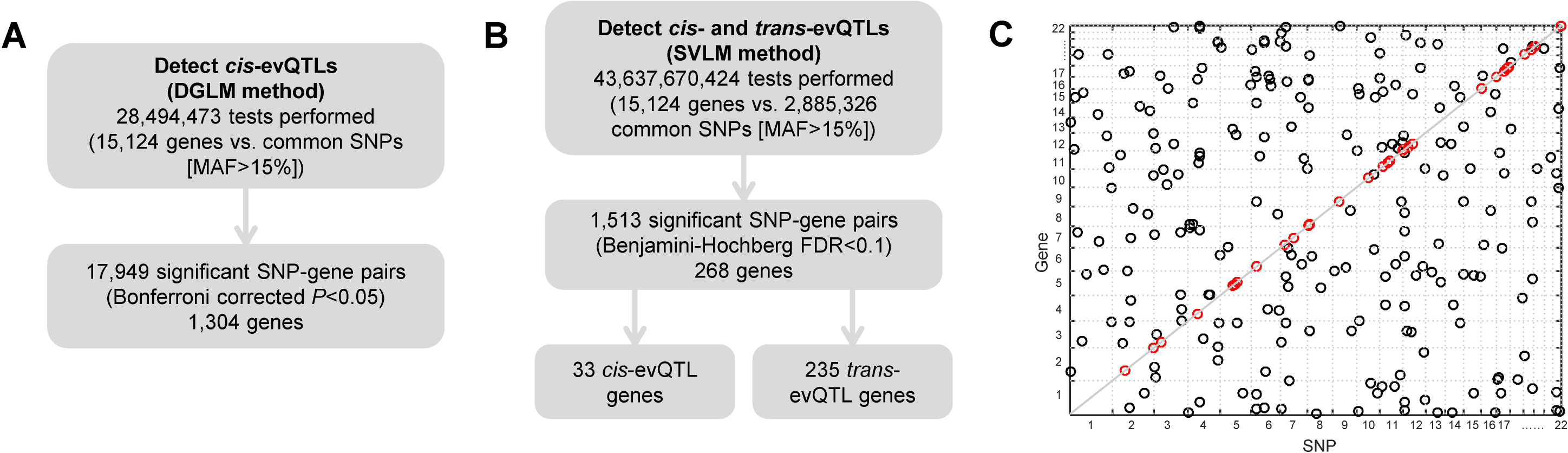
Overview of evQTL detections and the distribution of *cis*- and *trans*-evQTLs in autosomes of the human genome. (**A**) Flowchart of the identification of *cis*-evQTLs using the DGLM method. (**B**) Flowchart of the identification of *cis*- and *trans*-evQTLs using the SVLM method. (**C**) Distribution of *cis*- and *trans*-evQTLs identified using the SVLM method across autosomes.

### Epistasis contributes to increasing gene expression variability

Epistasis may cause an increase in the phenotypic variance of a population (10, 33). The effect of epistasis on gene expression variance can be interrogated using evQTLs (12). Here, we sought to identify “third-party” SNPs that epistatically interact with evSNPs and influence the expression of evQTL genes. For each evSNP, we identify the SNPs that influence the gene expression mean of a fraction of samples whose expression are more variable as defined by the evSNP. We call these SNPs the *partial eQTL SNPs* or peSNPs. They interact with evSNPs to increase the expression variance of the evQTL genes (9, 12). The procedure of peSNP identification is illustrated in **Supplementary Fig. S2**. Briefly, for a given evQTL (for example, an evQTL between gene ***G_expr_*** and ***SNP_ev_***), we extracted samples with the homozygous genotype associated with large expression variance. We called these L group samples. Accordingly, those related to small expression variance were named S group samples. Then, we conducted a genome-wide scan among the extracted L group samples to identify peSNPs (e.g., ***SNP_pe_***) that influence the expression of corresponding gene ***G_expr_***. The peSNPs are identified in the sub-sampled discovery panel, and their effect on gene expression is restricted to L group samples. ***SNP_ev_*** and ***SNP_pe_*** may be proximately co-localized on the same chromosome and partially associated as we showed previously (12). They may also be unlinked—for instance, located on different chromosomes—and interact with each other epistatically (9). We focused on the 268 evQTLs (33 *cis*- and 235 *trans*-acting) identified via SVLM. Among them, we identified that 73 evQTL genes harbor at least one significant interacting peSNP via simple linear regression test *P* < 10^-8^ on the L group samples (**Supplementary Table S3**). These results suggest that more than one-fourth of evQTLs are attributable to peSNPs interacting with evSNPs.

### Destabilizing evSNPs contribute to discordant gene expression between twin pairs

Here we present the destabilization model to explain the formation of evQTLs. The destabilization model concerns the destabilizing function of single SNPs that disrupt transcriptional machinery and cause variable gene expression. Unlike the epistatic evSNPs that take effect through genetic interactions with other SNPs (9, 12), destabilizing evSNPs increase gene expression variability without interacting with any other SNPs.

To show the destabilizing effects of SNPs, we re-visited the data derived from LCLs of a cohort of twin pairs of the TwinsUK project (34, 35), using monozygotic (MZ) twins to control for genetic background diversity among samples. In our previous study (12), we used the twin data for evQTL analysis and identified *cis*-evQTLs in 99 unique genes. Here, we first classified the 99 *cis*-evQTLs (between genes and the most significant SNPs) into 56 epistatic and 43 destabilizing evQTLs, based on whether an interacting SNP (i.e., peSNP) could be identified using the two-stage peQTL detection method described above. If no interacting SNP was detected, we associated the evQTL with the destabilization (rather than epistasis) model. Next, we extracted expression data for 139 pairs of MZ twins (34) (**Materials and Methods**). We classified MZ twin pairs with evSNP homozygous genotypes into either MZ-L or MZ-S groups based on whether the allele of the evSNP was associated with larger or smaller variance, relative to the alternative allele. We estimated the discordant gene expression between the two individuals of the same twin pairs using the *relative mean difference* (RMD), defined as the difference between two individuals’ gene expression values normalized by the mean. We then compared the average RMD between the MZ-L and MZ-S groups.

We used one destabilizing evQTL and one epistasis evQTL as examples to illustrate the difference between them in terms of the discordant gene expression between MZ-L and MZ-S groups. The destabilizing evQTL is between *TBKBP1* and rs1912483 (**Fig. 2A**, right), and the epistasis evQTL is between *PTER* and rs7913889 (**Fig. 2B**, right). The data points representing gene expression levels were grouped by the genotypes. Within each genotype category, data points from the same twin pair are displayed side-by-side. Every two individuals of the same MZ pair are linked by a gray line. The slope of the lines is an indicator of discordant gene expression between twin pairs. In the destabilizing evQTL example, the slopes between MZ twins with genotypes associated with large expression variance (i.e., MZ-L group) tend to be steeper than those with small expression variance (i.e., MZ-S group) (**Fig. 2A**, right). In contrast, in the epistasis evQTL example, the difference in slope skewness between MZ-L and MZ-S groups is less pronounced (**Fig. 2B**, right). We pooled RMD values from different twin pairs by MZ-L or MZ-S group and compared the distributions of RMD values between the two groups. For destabilizing evQTLs, the distributions of RMD values between MZ-L and MZ-S groups were significantly different (*P* = 1.3×10^-5^, **Fig. 2A**, left), with larger RMD values for the MZ-L group. For epistasis evQTLs, in contrast, this difference in RMD distribution was not detected between MZ-L and MZ-S groups (*P* = 0.052, **Fig. 2B**, left).

**Fig. 2.**
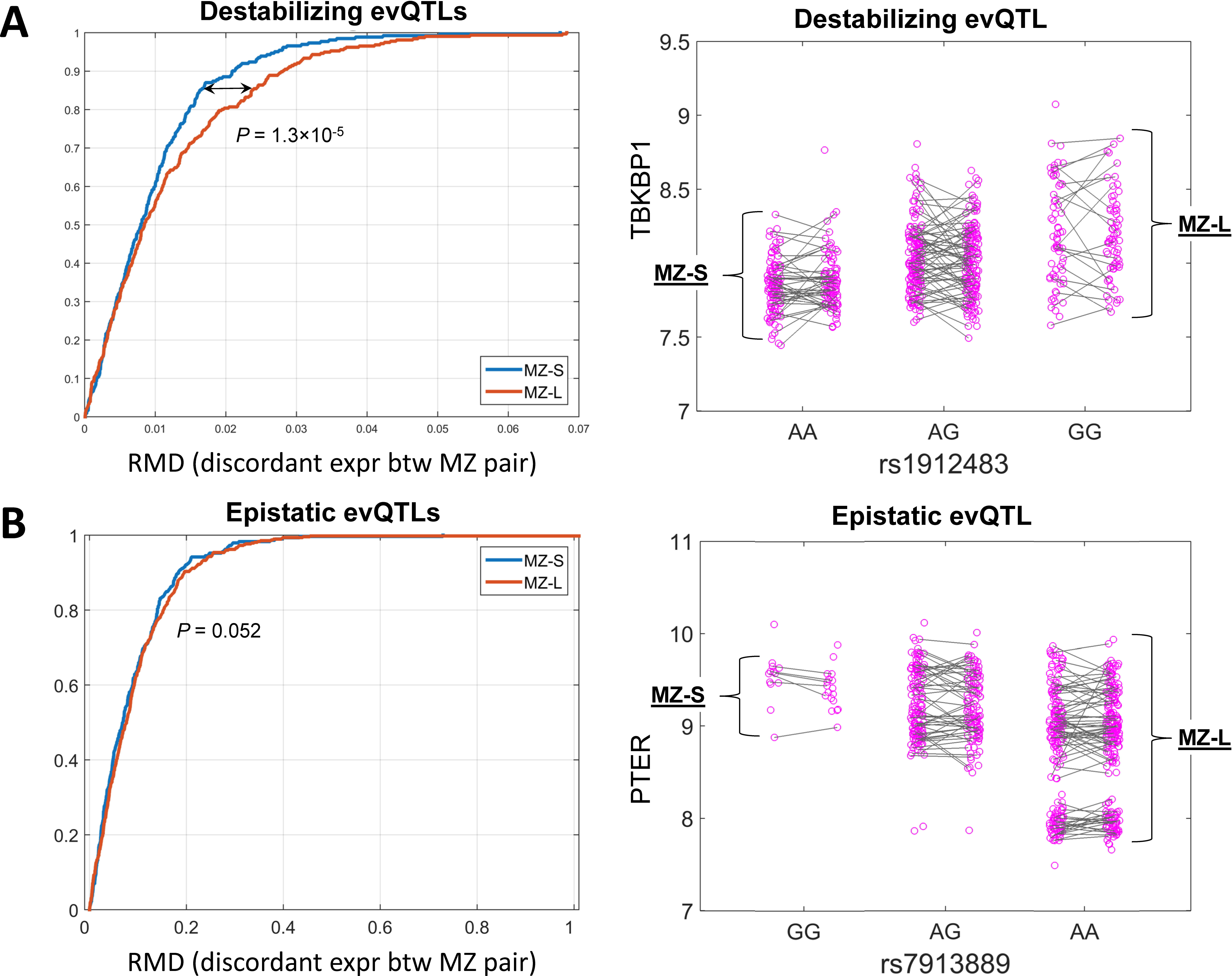
Dissection of epistatic and destabilizing effects of evQTLs using twins data. (A) The left panel shows the cumulative distribution function (CDF) curves of normalized discordant gene expression (measured as RMD) between MZ pairs in L and S (MZ-L and MZ-S) groups. The right panel shows an example of destabilizing evQTL between *TBKBP1* and rs1912483. The expression data points for each of two individuals from the same pairs of MZ twins are linked. Twin pairs are grouped as MZ-L and MZ-S based on whether the homozygous genotype at rs1912483 is associated with large or small gene expression variance. (B) Same as (A) but showing an example of epistatic evQTL between *PTER* and rs7913889.

### Destabilizing evSNPs may cause unstable gene expression of samples

Our destabilization model is based on the action of a single genetic variant that confers the destabilizing effect on gene expression. We hypothesized that different alleles of a destabilizing evSNP might be associated with different levels of gene expression stability in cell line samples. To test this hypothesis, we set out to estimate the time-course stability of gene expression by repeatedly measuring the gene expression level using qRT-PCR in each of the same cell lines multiple times. If our hypothesis is valid, then the stability of gene expression in samples with an evQTL genotype associated with larger variance should be lower than that in samples with an evQTL genotype associated with smaller variance.

We selected two destabilizing evQTLs for testing: *ATMIN*-rs1018804 and *BEND4*-rs7659929. *ATMIN* is an essential cofactor for the checkpoint kinase ATM, which transduces genomic stress signals to halt cell cycle progression and promote DNA repair (36). We picked two LCLs, HG00097 and HG00364, which have similar *ATMIN* expression levels. Both were derived from female individuals of European descent. At rs1018804, the CC genotype of HG00097 was associated with larger variance, while the AA genotype of HG00364 was associated with smaller variance. Thus, HG00097 and HG00364 belonged to L and S groups, respectively. We measured the evQTL gene expression level using qRT-PCR with three technical replicates at each of ten different sampling time points. The same assay was repeated three times independently. Our results showed that, under the same controlled experimental conditions, the variance of gene expression (i.e., the variance in ΔCt values) in HG00097 was greater than HG00364. The same trend was observed from all three biological replicates (**Fig. 3A**). In all three replicates, the difference was statistically significant (B-F test, *P* = 0.034, 0.019, and 0.0096, respectively).

**Fig. 3.**
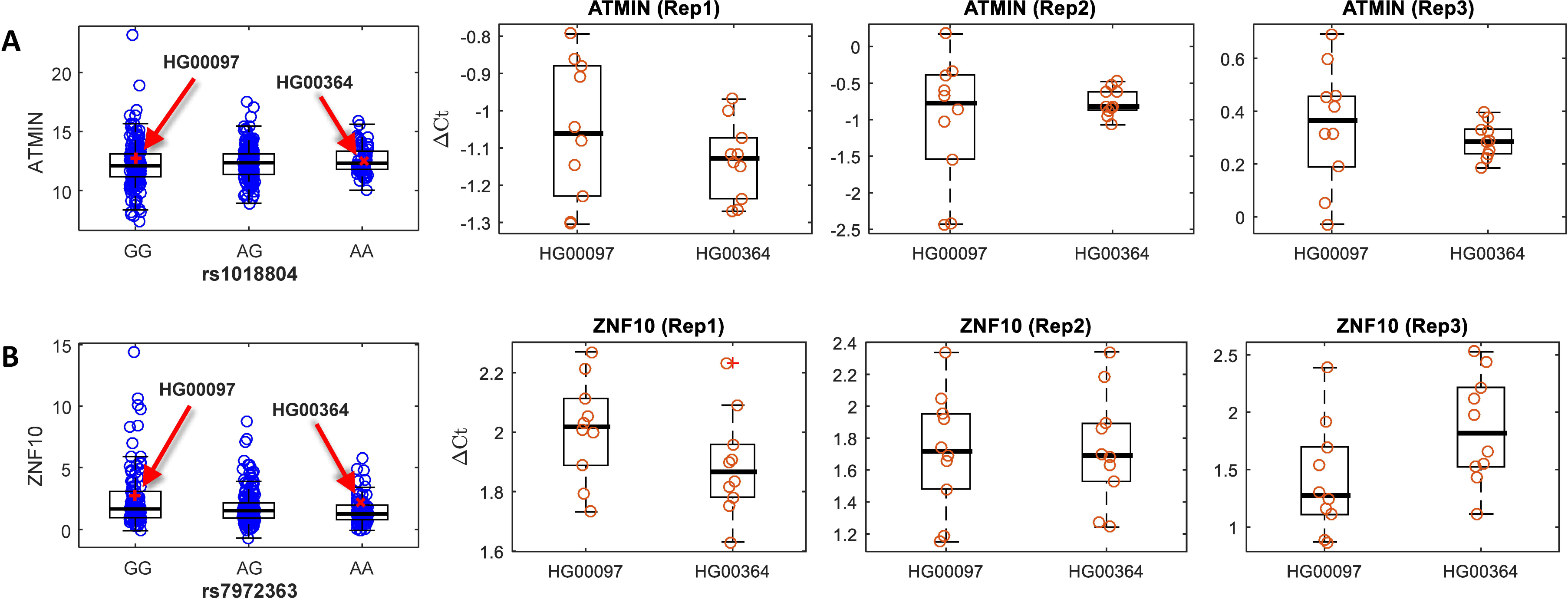
The negative correlation between gene expression variance and time-course stability presents in a destabilizing evQTL, but not in an epistasis evQTL, showing by the same cell lines. (A) The most left panel shows the gene expression levels of *ATMIN* among rs1018804 genotypes. Red arrows indicate the cell lines, HG00097 and HG00364. Right panels show the results of three replicates of the repetitive qRT-PCR analysis for *ATMIN* at ten different time points. At each time point of each biological replicate, three technical replicates are performed to obtain average ΔCt values, indicated by red circles. (B) Same as (A) but showing the results for the epistatic evQTL, *ZNF10*-rs7972363.

We repeated the experiment with two biological replicates on the same evQTL *ATMIN*-rs1018804 using a different pair of LCLs (NA12144 and NA12736 from L and S group, respectively) in place of HG00097 and HG00364. We obtained similar results showing a consistent pattern; that is, the gene expression in the cell line of L group was more variable than that of S group (**Supplementary Fig. S3**). Furthermore, we repeated the experiment on a different destabilizing evQTL (*BEND4*-rs7659929) with another pair of LCLs (NA12889 and NA18858). Again, we obtained the consistent pattern supporting the negative relationship between the population-level gene expression variance and the time-course stability of gene expression in single samples (**Supplementary Fig. S3**).

We further hypothesized that the negative relationship between gene expression variance and stability holds true exclusively for destabilizing evQTLs. Such a relationship should not be anticipated in epistasis evQTLs, because of the different modes of action through which the two kinds of evQTLs work. To test the hypothesis, we applied the same qRT-PCR strategy on an epistasis evQTL, *ZNF10*-rs7972363, using the same cell lines HG00097 and HG00364 (**Fig. 3B**). The AA genotype of HG00097 at rs7972363 is associated with larger variance while the GG genotype of HG00364 is associated with smaller variance. An epistasis evQTL, resulting from the interaction between SNPs rs1567910 and rs7972363, has been identified using the two-stage peQTL detection method. Samples with the AA genotype at rs7972363 can be further broken down by rs1567910 genotypes into three subgenotype groups associated with different levels of gene expression mean. As expected, the gene expression variance in ΔCt values was similar between HG00097 and HG00364 (**Fig. 3B**, B-F test, *P* = 0.96, 0.83, and 0.73, for the three replicates, respectively). Together, our results suggest that the level of gene expression stability—i.e., the time-course random fluctuation of gene expression—is associated with destabilizing evQTL, but not epistatic, SNPs.

### Differences in cell cycle status and alternative splicing do not account for the destabilizing function conferred by destabilizing evSNPs

Finally, we controlled for two additional confounding factors that might account for the increased gene expression variance associated with destabilizing evSNPs. The first is the cell cycle status of cell lines.

At a given sampling time, cells may consist various proportion of subpopulations at different cell cycle phases, which may contribute to the variance of expression observed in our measurement. To determine the potential impact of this factor, we performed a cell cycle study by flow cytometry with HG00097 and HG00364 at 36 h after incubation (**Materials and Methods**). The results showed no difference in the percentage of cells in G0/G1, S and G2/M phases between the two cell lines (**Supplementary Fig. S4**). The second confounding factor we considered is alternative splicing patterns. Different splicing patterns between cell lines might result in different gene-level expression measurements. We used the Integrative Genomics Viewer (37) to visualize the alternatively spliced mRNA of *ATMIN* and compared the pattern of splicing between HG00097 and HG00364, as well as that of *BEND4* between NA12889 and NA18858. In either case, we observed no difference in splicing patterns (**Supplementary Fig. S5**).

## Discussion

Emerging experimental and statistical techniques have enabled the rigorous analysis of phenotypic variability (15). Here, focusing on the variability of gene expression, we found that evQTLs are abundant and widespread across the human genome, which confirms our previous findings (11, 12). More specifically, we used the two-stage SVLM approach to the Geuvadis data of 345 EUR samples and identified 1,513 (505 *cis*- and 1,008 *trans*-) evQTLs at FDR of 10%. The numbers of evQTLs are different but largely comparable to those reported in a previous study by Brown et al. (9), in which 508 *cis*- and *trans*-evQTLs were identified at FDR of 5% with 765 LCLs from twin samples. The discrepancy in numbers of detected evQTLs between the two studies are attributable to differences in sample size, sampled populations, genotyping technique, and FDR cutoff.

Importantly, we discovered two distinct modes of action, through which common SNPs influence gene expression variance: epistasis and the destabilizing mutation. The epistasis model involves two or more variants that interact in a non-additive fashion (9, 38) or link to each other through incomplete linkage disequilibrium (12, 39). With this model, detecting increased phenotypic variance has been used to identify epistasis (10, 33, 40). The destabilization model, on the other hand, concerns the effect of single variants that work alone to destabilize phenotypic expression, pushing a proportion of individuals away from the robust optimum.

Dissecting epistatic and destabilizing effects in the context of evQTLs is technically challenging. Under the experimental conditions applied, we found that a quarter of evQTLs with detectable peSNPs could be explained by the epistasis model. However, a number of factors may cause the real fraction of epistatic evQTLs to be underestimated. These factors may include the low allele frequency and low effect size of either evSNPs or peSNPs, low power for detecting evSNPs or peSNPs due to the multiple testing burden, incomplete genotype information, and uncorrected environmental or technical effects. Indeed, in a previous study (12), we found that nearly half of evQTLs could be attributed to epistasis. Thus, caution should be taken when classifying any given evQTL as being explained by the destabilization model rather than the epistasis model, especially since our single epistatic variant (peSNP) detection ignores possible polygenic epistatic interactions. The absence of peSNP associated with an evQTL should not be considered as evidence of the absence of high-order epistatic interactions between SNPs, which are responsible for the formation of the evQTL.

The precise mechanisms underlying the destabilizing function of evSNPs are not completely understood. One possibility is that evSNPs act through genotype-environmental (GxE) interactions (41), e.g., the destabilizing mutations make the cellular transcriptional system more sensitive to the changing environment. To test this, we have taken the advantage of the identical genetic background of MZ twins and revealed that destabilizing evSNPs are associated with varying degrees of transcriptional stability. The increased discordant expression between MZ twins seems to support the GxE effect related to the destabilizing mechanism. Furthermore, destabilizing evSNPs are associated with increased transcriptional instability as shown by the repetitive qRT-PCR experiment. Nevertheless, future studies are needed to determine how the unstable expression of a gene in single samples could contribute to the gene expression variability of the population as a whole.

One of the underlying sources of the gene expression variability is the cell-to-cell variability in mRNA levels, typically referred to as stochastic noise (6, 42–45). Recently developed single-cell technologies (46) have enabled the measurement of the cell-to-cell variability in gene expression (6, 43–45). Destabilizing mutations may include those that affect transcription factor binding (6), the balance between promoter activation and transcription (44), the burst frequency and burst size (45), and the genetic redundancy of transcriptional networks (43). Although we did not evaluate the destabilizing function conferred by evSNPs at the single-cell level in this study, we hypothesize that decreased cell-to-cell variability in gene expression, i.e., more homogeneous expression across single cells, may result in less stable gene expression of samples. This is because the highly coordinated regulation of transcription is likely to drive all cells to be responsive to the same regulatory signal, then when the signal changes, the new signal will drive the whole population of cells to another level away from the previously established level. Hence, the sample shows a reduced stability of gene expression. Conversely, a high level of transcriptional heterogeneity across single cells may result in a more stable gene expression, as the cell-to-cell stochastic noise would cancel out when samples of many cells are evaluated.

Despite many unknowns and the need for extra effort to reveal the precise mechanisms of evSNP function, we anticipate our findings have many implications. For example, given that regulatory variations play critical roles in many human diseases (47), understanding how genetic variation contributes to increasing gene expression variability will facilitate the identification of disease-related variants. This is especially true when gene expression heterogeneity characterizes traits or diseases such as aging (48–50) and cancer (51). For many diseases that display a high degree of phenotypic heterogeneity among patients, we may consider that the increased phenotypic variability is due to variability-controlling mutations (such as evSNPs). The improved insight of how these mutations influence the variability may bring us closer to causal mutations underlying individuals’ predispositions to disease. This strategy, in combination with other methods for estimating the impact of rare mutations, such as the aberrant gene expression analysis we have developed (52), would be valuable in the pursuit of personalized medicine. Furthermore, we suggest that variability-controlling mutations could be potential targets for genomic editing or drug development.

In conclusion, we described a new groups evSNPs that may act to destabilize the gene transcription, which is distinct from the previously reported epistatic SNPs. Destabilization SNPs act independently from other SNPs, whereas epistatic evQTLs can interact with each other to influence gene expression variability. Collectively, SNPs of both action models assist shaping the variability of gene expression in a population and contribute to the formation of evQTLs. Our findings provide unique entry points to follow-up mechanistic analyses, which may open up new, variability-centered research avenues for mapping complex traits. Methods derived from such a new analytical framework may allow us to identify novel causal loci, which would otherwise be missed by traditional mean-focused methods.

## Materials and Methods

### Gene expression and genotype data for evQTL analysis

The Geuvadis RNA-seq data were downloaded from the EBI ArrayExpress website using accession EGEUV-1 (28). The data matrix contained the expression levels of Gencode (v12)-annotated genes in 462 unique LCL samples. The data had been quantile normalized and further processed using the method of probabilistic estimation of expression residuals (PEER) (53). From the downloaded data matrix, we extracted the expression values of autosomal protein-coding genes in 345 EUR samples, whose genotype data is available from the website of the 1,000 Genomes Project (29). Based on the result of the principal component analysis, we excluded 19 samples whose global expression profiles deviated the most from those of the rest of the samples. The final expression matrix contained the data of 15,124 protein-coding genes for 326 EUR samples. We also obtained the genotype and expression data from the TwinsUK project (34, 35). The expression data from 139 pairs of MZ twins (34) were extracted as previously described (12).

### Identification of evQTLs

We adopted the DGLM method to test for inequality in expression variances and measure the contribution of genetic variants to expression heteroscedasticity. We considered the following model: *y_i_* = *μ* + *x_i_β* + *g_i_α* + *∈_i_*, *∈_i_* ∼*N*(0, σ^2^ exp(*g_i_θ*)), where *y_i_* indicates a gene expression trait of individual *i*, *g_i_* is the genotype at the given SNP (encoded as 0, 1, or 2 for homozygous rare, heterozygous and homozygous common alleles, respectively), □_*i*_ is the residual with variance σ^2^, and *θ* is the corresponding vector of coefficients of genotype *g_i_* on the residual variance. Ages of subjects and the batch of data collection were modeled as covariates *x_i_*. With this full model, both mean and variance of expression *y_i_* were controlled by SNP genotype *g_i_*. We also used the SVLM procedure (31) to detect evQTLs. The SVLM method consists of two steps. First, a regression analysis is applied where the gene expression trait is adjusted for a possible SNP effect and the effect of other covariates is also regressed out. Second, another regression analysis is applied to the squared values of residuals obtained from the first stage, using the SNP as the predictor to capture the effect of the SNP on the expression residuals. Simulation-based analysis was performed to estimate the power of the two evQTL detection methods. Briefly, a population of 10,000 individuals with a hypothetical evSNP with a minor allele frequency (MAF) of 0.4 was generated. Following the Hardy–Weinberg equilibrium, the 10,000 simulated individuals were randomly assigned into one of three genotype groups indicated with 0, 1, 2 for the number of minor allele of the evSNP. The gene expression variance was set as 1.0, 2.0 and 4.0, respectively, and the mean was set as zero the three genotype groups. Two evQTL detection methods DGLM and SVLM were applied to the same simulated population and each was repeated for 1,000 times to estimate the rate of positive detection, i.e., the power of the method. The whole analysis was repeated with different sample sizes ranging from 300 to 1,000 with a step of 100 to establish the power as a function of varying sample size (**Supplementary Fig. S1**).

### Identification of peSNPs that interact with evSNPs

We used a two-stage procedure to identify peSNPs that interact with evQTLs. We first partitioned individuals into L and S groups according to whether genotypes of the evSNP are associated with large (L) and small (S) variance of gene expression. Then we scanned genome-wide SNPs. For each SNP, the eQTL analysis by linear regression model was conducted among individuals of the L group. For each top SNP with high genotype heterozygosity difference, a linear regression (54) was performed on the SNP’s genotypes and gene expression. The most significant SNPs were retained after applying an arbitrary *P*-value = 0.0005 as cutoff and were reported as candidate interacting SNPs.

### Estimation of time-course stability of gene expression using repetitive qRTPCR assay

LCLs were purchased from the Coriell Institute (https://catalog.coriell.org/). Cells were maintained in Roswell Park Memorial Institute Medium 1640 supplemented with 2mM L-glutamine and 15% FBS (Seradigm) at 37°C in a humidified atmosphere containing 5% CO_2_ (v/v). For the time course experiment, cell lines were seeded at 1×10^6^ cells per 10 cm dish and then incubated in the culture medium. Cell lines were screened using the MycoFluor mycoplasma detection kit (Invitrogen) to ensure their mycoplasma free condition. Cells were collected at ten different time points from 12 to 72 h after seeding. Total RNA was extracted using Trizol reagent (Invitrogen). RNase-free DNase (Ambion) was used to remove trace amount of potential genomic DNA contamination. RNA purity and concentration were determined using a Nanodrop ND-100 Spectrophotometer. The concentrations of total RNA were adjusted to 100 μg/ml. Real-time PCR assays were performed using iTaq Universal SYBR Green One-Step Kit (Bio-Rad Laboratories) with primers shown in **Supplementary Table S4**. Template total RNA was reverse transcribed and amplified in a Bio-Rad CFX96 Real-Time PCR Detection System (Bio-Rad Laboratories) in 20-μl reaction mixtures containing 10 μl of iTaq universal SYBR Green reaction mix (2×), 0.25 μl of iScript reverse transcriptase, 2 μl of 100 nM of forward and reverse primers mix, 1 μl of total RNA template, and 6.75 μl of nuclease-free water, at 50°C for 10 min, 95°C for 1 min, followed by 40 cycles of 95°C for 10 s and 58°C for 30 s. Melting curves were measured from 65°C to 95°C with 0.5°C of increment. The average expression of two housekeeping genes, *CHMP2A* and *C1orf43*, was used for normalization. The choice of using these two genes as reference was based on a recent RNA-seq study for constantly expressed human genes (55). The expression stability of the two genes was further confirmed by using geNorm and NormFinder programs (56, 57).

### Flow cytometric analysis of cells in different phases of the cell cycle

The proportion of cells at different cell cycle statuses was determined using flow cytometry analysis, based on the measurement of the DNA content of nuclei labeled with propidium iodide (PI). Cells were harvested at 24, 36, 48, 60, and 72 h after plating. Cells were resuspended at a concentration of 1×10^6^/ml in cold PBS. After 1 ml of ice-cold 100% ethanol had been added dropwise, the cells were fixed at 4°C for at least 16 hours. Fixed cells were pelleted, resuspended in 1ml of PI staining solution (50 mg/ml propidium iodide, 100 units/ml RNase A in PBS) for at least 1 hour at room temperature and analyzed on an FACS flow cytometer (BD Biosciences). By using red propidium-DNA fluorescence, 30,000 events were acquired. The percentage of cells in G0/G1, S and G2/M phases of the cell cycle was calculated using Flowjo v10 (Tree Star).

## Acknowledgements

We thank Candice Brinkmeyer-Langford for critically reading and editing this manuscript. We thank Guan Zhu for valuable discussion, Jinting Guan for help with computer simulations, Srikanth Kanameni for help with flow cytometry, and Vijayanagaram Venkatraj for help with cell line preparation. We acknowledge the Texas A&M Institute for Genome Sciences and Society (TIGSS) for providing computational resources. The TwinsUK study was funded by the Wellcome Trust and the European Community’s Seventh Framework Programme (FP7/2007–2013). The study also received support from the National Institute for Health Research (NIHR)’s Clinical Research Facility at Guy’s and St. Thomas’ National Health Service (NHS) Foundation Trust and NIHR’s Biomedical Research Centre based at Guy’s and St. Thomas’ NHS Foundation Trust and King’s College London. SNP genotyping was performed by the Wellcome Trust Sanger Institute and the National Eye Institute via National Institutes of Health/Computerized Infectious Disease Reporting (CIDR).

## Conflict of Interest Statement

None.

## Supplementary Figure Legends

**Supplementary Fig. S1.** Comparison of statistical power of two evQTL detection methods: DGLM and SVLM, using computer simulations with different sample sizes. For simulations, a population of 10,000 individuals was generated, and the MAF of an evSNP was set to 0.4. The genotypes of SNP were encoded to 0, 1, 2 for homozygous minor, heterozygous, and homozygous major alleles, respectively. The gene expression of each genotype was generated from a normal distribution with the same mean but different variances, 1.0, 2.0, and 4.0, respectively. Before testing a method, the population was subsampled to the designated sample size, ranging from 300 to 1,000. For each sample size, the tested method was applied to the subsamples. The whole procedure was repeated 1,000 times, and the power was computed as the ratio of the times of *P*-value being smaller than 5×10^-5^ (i.e., 0.05/1000).

**Supplementary Fig. S2.** Schematic illustration of the method for identifying peQTLs. After the identification of evQTL, the peQTL method involves two steps: (1) extraction of homozygous individuals whose genotype of the evSNP is associated with increased expression variability, and (2) identification of the eQTL between the gene and third-party variant among extracted individuals.

**Supplementary Fig. S3.** The anti-correlation between the levels of gene expression variance and the time-course measurements of transcriptional stability presents in two additional destabilizing evQTLs. (A) Destabilizing evQTL *ATMIN*-rs1018804 gene expression in the cell line pair NA12144 and NA12763. The most left panel shows the distribution of gene expression levels of *ATMIN* among three different genotypes defined by two alleles of rs1018804. Red arrows indicate the expression levels of NA12144 and NA12736 and their genotypes. Right panels show the results of two biological replicates of repetitive qRT-PCR analysis for *ATMIN* at five different time points at 24, 36, 48, 60, and 72 h. At each time point of each biological replicate, three technical replicates were performed to obtain ΔCt values, and the average is presented by the red circle. (B) Same as (A) but showing the results of evQTL *BEND4*-rs7659929 using cell line pair NA12889 and NA18858.

**Supplementary Fig. S4.** Cell cycle analysis to determine the relative abundance of cells in different phases. (**A**) Representative flow cytometric dot plots. (**B**) Representative histograms obtained using TUNEL assay. (**C**) Relative frequencies of cells in G1, S, and G2 phases. (**D**) Principal component analysis of cell cycle profiles. (**E**) Relative frequencies of cells in different phases of HG00097 (red) and HG00364 (blue).

**Supplementary Fig. S5.** The IGV view of RNA-seq read alignments and the sashimi plot of mRNA splicing patterns of evQTL genes in different cell lines. (A) IGV view of RNA-seq read alignment of *ATMIN* in HG00097 and HG00364. (B) Sashimi plots of *ATMIN* mRNA in HG00097 and HG00364. (C) IGV view of RNA-seq read alignment of *BEND4* in NA12889 and NA18858. (D) Sashimi plots of *BEND4* mRNA in NA12889 and NA18858.

## Supplementary Table Legends

**Supplementary Table S1.** The list of *cis*-evQTLs identified using the DGLM method.

**Supplementary Table S2.** The list of *cis*- and *trans*-evQTLs identified using the SVLM method.

**Supplementary Table S3.** The list of peQTLs and the corresponding evQTLs.

**Supplementary Table S4.** Sequences of primers used in the qRT-PCR assay.

## References

1 Lynch, M. and Walsh, B. (1998) Genetics and analysis of quantitative traits. Sinauer, Sunderland, Mass.

2 Yang, J., Loos, R.J., Powell, J.E., Medland, S.E., Speliotes, E.K., Chasman, D.I., Rose, L.M., Thorleifsson, G., Steinthorsdottir, V., Magi, R. et al. (2012) FTO genotype is associated with phenotypic variability of body mass index. Nature, 490, 267–272.

3 Jimenez-Gomez, J.M., Corwin, J.A., Joseph, B., Maloof, J.N. and Kliebenstein, D.J. (2011) Genomic analysis of QTLs and genes altering natural variation in stochastic noise. PLoS genetics, 7, e1002295.

4 Ansel, J., Bottin, H., Rodriguez-Beltran, C., Damon, C., Nagarajan, M., Fehrmann, S., Francois, J. and Yvert, G. (2008) Cell-to-cell stochastic variation in gene expression is a complex genetic trait. PLoS genetics, 4, e1000049.

5 Ronnegard, L. and Valdar, W. (2011) Detecting major genetic loci controlling phenotypic variability in experimental crosses. Genetics, 188, 435–447.

6 Metzger, B.P., Yuan, D.C., Gruber, J.D., Duveau, F. and Wittkopp, P.J. (2015) Selection on noise constrains variation in a eukaryotic promoter. Nature, 521, 344–347.

7 Shen, X., Pettersson, M., Ronnegard, L. and Carlborg, O. (2012) Inheritance beyond plain heritability: variance-controlling genes in Arabidopsis thaliana. PLoS genetics, 8, e1002839.

8 Ayroles, J.F., Buchanan, S.M., O’Leary, C., Skutt-Kakaria, K., Grenier, J.K., Clark, A.G., Hartl, D.L. and de Bivort, B.L. (2015) Behavioral idiosyncrasy reveals genetic control of phenotypic variability. Proc Natl Acad Sci U S A, 112, 6706–6711.

9 Brown, A.A., Buil, A., Vinuela, A., Lappalainen, T., Zheng, H.F., Richards, J.B., Small, K.S., Spector, T.D., Dermitzakis, E.T. and Durbin, R. (2014) Genetic interactions affecting human gene expression identified by variance association mapping. Elife, 3, e01381.

10 Pare, G., Cook, N.R., Ridker, P.M. and Chasman, D.I. (2010) On the use of variance per genotype as a tool to identify quantitative trait interaction effects: a report from the Women’s Genome Health Study. PLoS genetics, 6, e1000981.

11 Hulse, A.M. and Cai, J.J. (2013) Genetic variants contribute to gene expression variability in humans. Genetics, 193, 95–108.

12 Wang, G., Yang, E., Brinkmeyer-Langford, C.L. and Cai, J.J. (2014) Additive, epistatic, and environmental effects through the lens of expression variability QTL in a twin cohort. Genetics, 196, 413–425.

13 Fehrmann, S., Bottin-Duplus, H., Leonidou, A., Mollereau, E., Barthelaix, A., Wei, W., Steinmetz, L.M. and Yvert, G. (2013) Natural sequence variants of yeast environmental sensors confer cell-to-cell expression variability. Mol Syst Biol, 9, 695.

14 Chuffart, F., Richard, M., Jost, D., Burny, C., Duplus-Bottin, H., Ohya, Y. and Yvert, G. (2016) Exploiting Single-Cell Quantitative Data to Map Genetic Variants Having Probabilistic Effects. PLoS genetics, 12, e1006213.

15 Geiler-Samerotte, K.A., Bauer, C.R., Li, S., Ziv, N., Gresham, D. and Siegal, M.L. (2013) The details in the distributions: why and how to study phenotypic variability. Curr Opin Biotechnol, 24, 752–759.

16 Gibson, G. (2009) Decanalization and the origin of complex disease. Nature reviews. Genetics, 10, 134–140.

17 Queitsch, C., Sangster, T.A. and Lindquist, S. (2002) Hsp90 as a capacitor of phenotypic variation. Nature, 417, 618–624.

18 Gibson, G. and Wagner, G. (2000) Canalization in evolutionary genetics: a stabilizing theory? Bioessays, 22, 372–380.

19 Wolf, L., Silander, O.K. and van Nimwegen, E. (2015) Expression noise facilitates the evolution of gene regulation. Elife, 4.

20 Zhang, Z., Qian, W. and Zhang, J. (2009) Positive selection for elevated gene expression noise in yeast. Mol Syst Biol, 5, 299.

21 Acar, M., Mettetal, J.T. and van Oudenaarden, A. (2008) Stochastic switching as a survival strategy in fluctuating environments. Nat Genet, 40, 471–475.

22 Kaern, M., Elston, T.C., Blake, W.J. and Collins, J.J. (2005) Stochasticity in gene expression: from theories to phenotypes. Nature reviews. Genetics, 6, 451–464.

23 Hill, W.G. and Zhang, X.S. (2004) Effects on phenotypic variability of directional selection arising through genetic differences in residual variability. Genet Res, 83, 121–132.

24 Hill, W.G. and Mulder, H.A. (2010) Genetic analysis of environmental variation. Genet Res (Camb), 92, 381–395.

25 Mar, J.C., Matigian, N.A., Mackay-Sim, A., Mellick, G.D., Sue, C.M., Silburn, P.A., McGrath, J.J., Quackenbush, J. and Wells, C.A. (2011) Variance of gene expression identifies altered network constraints in neurological disease. PLoS genetics, 7, e1002207.

26 Ho, J.W., Stefani, M., dos Remedios, C.G. and Charleston, M.A. (2008) Differential variability analysis of gene expression and its application to human diseases. Bioinformatics, 24, i390–398.

27 Campbell, M.G., Kohane, I.S. and Kong, S.W. (2013) Pathway-based outlier method reveals heterogeneous genomic structure of autism in blood transcriptome. BMC Med Genomics, 6, 34.

28 Lappalainen, T., Sammeth, M., Friedlander, M.R., t Hoen, P.A., Monlong, J., Rivas, M.A., Gonzalez-Porta, M., Kurbatova, N., Griebel, T., Ferreira, P.G. et al. (2013) Transcriptome and genome sequencing uncovers functional variation in humans. Nature, 501, 506–511.

29 Abecasis, G.R., Auton, A., Brooks, L.D., DePristo, M.A., Durbin, R.M., Handsaker, R.E., Kang, H.M., Marth, G.T. and McVean, G.A. (2012) An integrated map of genetic variation from 1,092 human genomes. Nature, 491, 56–65.

30 Verbyla, A.P. and Smyth, G.K. (1998) Double generalized linear models: approximate residual maximum likelihood and diagnostics. Research Report, 1–15.

31 Struchalin, M.V., Amin, N., Eilers, P.H., van Duijn, C.M. and Aulchenko, Y.S. (2012) An R package “VariABEL” for genome-wide searching of potentially interacting loci by testing genotypic variance heterogeneity. BMC Genet, 13, 4.

32 Benjamini, Y. and Hochberg, Y. (1995) Controlling the False Discovery Rate - a Practical and Powerful Approach to Multiple Testing. J Roy Stat Soc B Met, 57, 289–300.

33 Struchalin, M.V., Dehghan, A., Witteman, J.C., van Duijn, C. and Aulchenko, Y.S. (2010) Variance heterogeneity analysis for detection of potentially interacting genetic loci: method and its limitations. BMC Genet, 11, 92.

34 Grundberg, E., Small, K.S., Hedman, A.K., Nica, A.C., Buil, A., Keildson, S., Bell, J.T., Yang, T.P., Meduri, E., Barrett, A. et al. (2012) Mapping cis- and trans-regulatory effects across multiple tissues in twins. Nat Genet, 44, 1084–1089.

35 Moayyeri, A., Hammond, C.J., Hart, D.J. and Spector, T.D. (2013) The UK Adult Twin Registry (TwinsUK Resource). Twin Res Hum Genet, 16, 144–149.

36 Kanu, N. and Behrens, A. (2008) ATMINistrating ATM signalling: regulation of ATM by ATMIN. Cell Cycle, 7, 3483–3486.

37 Thorvaldsdottir, H., Robinson, J.T. and Mesirov, J.P. (2013) Integrative Genomics Viewer (IGV): high-performance genomics data visualization and exploration. Brief Bioinform, 14, 178–192.

38 Hemani, G., Shakhbazov, K., Westra, H.J., Esko, T., Henders, A.K., McRae, A.F., Yang, J., Gibson, G., Martin, N.G., Metspalu, A. et al. (2014) Detection and replication of epistasis influencing transcription in humans. Nature, 508, 249–253.

39 Wood, A.R., Tuke, M.A., Nalls, M.A., Hernandez, D.G., Bandinelli, S., Singleton, A.B., Melzer, D., Ferrucci, L., Frayling, T.M. and Weedon, M.N. (2014) Another explanation for apparent epistasis. Nature, 514, E3–5.

40 Daye, Z.J., Chen, J. and Li, H. (2012) High-Dimensional Heteroscedastic Regression with an Application to eQTL Data Analysis. Biometrics, 68, 316–326.

41 Buil, A., Brown, A.A., Lappalainen, T., Vinuela, A., Davies, M.N., Zheng, H.F., Richards, J.B., Glass, D., Small, K.S., Durbin, R. et al. (2015) Gene-gene and gene-environment interactions detected by transcriptome sequence analysis in twins. Nat Genet, 47, 88–91.

42 Elowitz, M.B., Levine, A.J., Siggia, E.D. and Swain, P.S. (2002) Stochastic gene expression in a single cell. Science, 297, 1183–1186.

43 Raj, A., Rifkin, S.A., Andersen, E. and van Oudenaarden, A. (2010) Variability in gene expression underlies incomplete penetrance. Nature, 463, 913–918.

44 Raser, J.M. and O’Shea, E.K. (2004) Control of stochasticity in eukaryotic gene expression. Science, 304, 1811–1814.

45 Hornung, G., Bar-Ziv, R., Rosin, D., Tokuriki, N., Tawfik, D.S., Oren, M. and Barkai, N. (2012) Noise-mean relationship in mutated promoters. Genome Res, 22, 2409–2417.

46 Dey, S.S., Foley, J.E., Limsirichai, P., Schaffer, D.V. and Arkin, A.P. (2015) Orthogonal control of expression mean and variance by epigenetic features at different genomic loci. Mol Syst Biol, 11, 806.

47 Albert, F.W. and Kruglyak, L. (2015) The role of regulatory variation in complex traits and disease. Nature reviews. Genetics, 16, 197–212.

48 Bahar, R., Hartmann, C.H., Rodriguez, K.A., Denny, A.D., Busuttil, R.A., Dolle, M.E., Calder, R.B., Chisholm, G.B., Pollock, B.H., Klein, C.A. et al. (2006) Increased cell-to-cell variation in gene expression in ageing mouse heart. Nature, 441, 1011–1014.

49 Vinuela, A., Brown, A.A., Buil, A., Tsai, P.-C., Davies, M.N., Bell, J.T., Dermitzakis, E., Spector, T. and Small, K. (2016) Age-dependent changes in mean and variance of gene expression across tissues in a twin cohort. bioRxiv, doi: 10.1101/063883.

50 Brinkmeyer-Langford, C.L., Guan, J., Ji, G. and Cai, J.J. (2016) Aging Shapes the Population-Mean and -Dispersion of Gene Expression in Human Brains. Frontiers in Aging Neuroscience, 8, doi: 10.3389/fnagi.2016.00183.

51 Ecker, S., Pancaldi, V., Rico, D. and Valencia, A. (2015) Higher gene expression variability in the more aggressive subtype of chronic lymphocytic leukemia. Genome Med, 7, 8.

52 Zeng, Y., Wang, G., Yang, E., Ji, G., Brinkmeyer-Langford, C.L. and Cai, J.J. (2015) Aberrant gene expression in humans. PLoS genetics, 11, e1004942.

53 Stegle, O., Parts, L., Durbin, R. and Winn, J. (2010) A Bayesian framework to account for complex non-genetic factors in gene expression levels greatly increases power in eQTL studies. PLoS Comput Biol, 6, e1000770.

54 Stranger, B.E., Forrest, M.S., Clark, A.G., Minichiello, M.J., Deutsch, S., Lyle, R., Hunt, S., Kahl, B., Antonarakis, S.E., Tavare, S. et al. (2005) Genome-wide associations of gene expression variation in humans. PLoS genetics, 1, e78.

55 Eisenberg, E. and Levanon, E.Y. (2013) Human housekeeping genes, revisited. Trends Genet, 29, 569–574.

56 Vandesompele, J., De Preter, K., Pattyn, F., Poppe, B., Van Roy, N., De Paepe, A. and Speleman, F. (2002) Accurate normalization of real-time quantitative RT-PCR data by geometric averaging of multiple internal control genes. Genome Biol, 3, RESEARCH0034.

57 Andersen, C.L., Jensen, J.L. and Orntoft, T.F. (2004) Normalization of real-time quantitative reverse transcription-PCR data: a model-based variance estimation approach to identify genes suited for normalization, applied to bladder and colon cancer data sets. Cancer Res, 64, 5245–5250.

